# Recall tempo of Hebbian sequences depends on the interplay of Hebbian kernel with tutor signal timing

**DOI:** 10.1101/2023.06.07.542926

**Authors:** Matthew Farrell, Cengiz Pehlevan

## Abstract

Understanding how neural circuits generate sequential activity is a longstanding challenge. While foundational theoretical models have shown how sequences can be stored as memories with Hebbian plasticity rules, these models considered only a narrow range of Hebbian rules. Here we introduce a model for arbitrary Hebbian plasticity rules, capturing the diversity of spike-timing-dependent synaptic plasticity seen in experiments, and show how the choice of these rules and of neural activity patterns influences sequence memory formation and retrieval. In particular, we derive a general theory that predicts the speed of sequence replay. This theory lays a foundation for explaining how cortical tutor signals might give rise to motor actions that eventually become “automatic”. Our theory also captures the impact of changing the speed of the tutor signal. Beyond shedding light on biological circuits, this theory has relevance in artificial intelligence by laying a foundation for frameworks whereby slow and computationally expensive deliberation can be stored as memories and eventually replaced by inexpensive recall.

## Introduction

An important class of animal behaviors are those that are consolidated into “automatic”, well-practiced sequential routines (1–9). An analogy is the process of learning a tennis serve, which starts as a slow and deliberate process, but eventually becomes nearly muscle-memory. Importantly, the behavior can be thought of as a sequence that, once initialized in a starting state, progresses in a highly stereotyped, automatic fashion. From this perspective, the behavior can be thought of as a stored sequence memory that is recalled by an initial prompt.

For such sequence memories, the timing of the sub-elements of these sequences and overall tempo are essential components (10–16). It is therefore important to understand how temporally specific sequences can be learned, memorized, and recalled.

To answer this question, we need to look at the mechanisms of learning and generation of sequential behavior. One common perspective holds that these behaviors are generated by stereotyped sequential neural activity (17–21). Further, several experiments showed a tight correlation between neural activity and timing of behavior (22–30). This suggests that automatic sequential behavior may at least in part be “stored” in the synapses of a neural circuit, such that the behavior can be generated by setting the network to a state that corresponds to the first point in the sequence.

Despite the importance of these topics to understanding brain function, it is unclear what actually determines the tempo of sequence generation in neural circuits, and how this is connected to the learning process. Here we hypothesize that the tempo arises from an interaction between the temporal dependence of the synaptic learning rules and the temporal structure of the network activity when learning the sequential behavior. We refer to the mechanism that sets network activity during sequence learning as the “tutor signal”; this tutor signal may come from a higher-order brain area such as motor cortex, or from minimally processed sensory inputs.

This perspective is embodied by models featuring temporally asymmetric Hebbian (TAH) learning rules. Despite the long and illustrious legacy of this class of models (31–41), previous studies were seriously limited by the very narrow class of learning rules that these preexisting models can describe, and the fact that these learning rules do not correspond with those observed in the brain. In this work we extend the basic TAH models previously considered to a much richer class of models. Not only do we derive a theory that describes the essential elements of sequence dynamics, including tempo and noise robustness, for learning rules currently seen in the brain; our theory describes a wide class of TAH learning rules that gives rise to sequential dynamics, covering the case of new learning rules that may be discovered in biology in the future.

TAH network models are built upon Hebbian learning rules, wherein the network state is set by the tutor to a sequence of activity patterns one after another. These activity patterns correspond to moments in behavioral sequences, and the Hebbian learning rules store the structure of these sequences in the network as a memory. The sequences can then be recalled by setting the network to the first state in a desired sequence, resulting in the execution of the now “automatic”, well-practiced sequential behavior. These models can be seen as a generalization of Hopfield networks (42) to storing sequences rather than static patterns; therefore, many of the motivations and rationale behind the Hopfield model is applicable here, such as the ability to “complete” a sequence based on partial inputs (see Discussion for more in-depth interpretations of this model).

While our theory is very general, we demonstrate its applicability and utility by focusing on two fundamental and biologically-relevant cases: (1) TAH learning rules with fast timescales relative to the tutor signal and (2) Double-sided exponential TAH learning rules as are commonly seen in biology (43). In both cases we find useful principles that are interpretable in the context of biology. For example, weight structures (and TAH learning rules) can be adjusted such that the tempo remains the same, but robustness to noise increases. We also show how networks with the same qualitative dynamical properties can arise from very different weight structure. For double-sided exponential TAH kernels, we show that the tempo of sequence recall depends linearly on the speed of the tutor signal, with a slope that depends in a simple way on the parameters of the kernel. While we focus here on applications to neuroscience, considering its generality our analysis may also be of interest to theorists in other domains studying dynamical systems with asymmetric weight structures.

## Results

### Sequence model definition

The sequence is stored and produced by a recurrently connected network of *N* neurons. The connectivity is given by a recurrent weight matrix *W*, where *W*_*ij*_ is the connectivity strength of neuron *j* to neuron *i*. The dynamics of the network are then given by

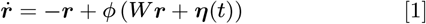

where ***η***(*t*) is white noise (⟨ *η*_*i*_(*t*)⟩ = 0 and ⟨ *η*_*i*_(*t*)*η*_*i*_(*t*^′^)⟩ = *ρ*^2^*δ*(*t*− *t*^′^)) and *ϕ* is a sigmoidal nonlinearity (see Appendix 1 for more details). Note that it is common to add a timescale factor *τ* which multiplies the term 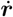 and 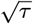 that multiplies *η*(*t*)), which have the effect of rescaling the timescale of the dynamics. Since this effect is relatively simple to understand, we simply take *τ* = 1 and focus our attention on other attributes.

The sequence is stored in this network by a tutor signal (Fig. 1a). Given a sequence of length *P*, this signal sets the state of the network sequentially to an ordered set of *P* patterns 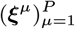. These *P* patterns represent *P* neural states that correspond to the states making up the sequential behavior. The recurrent network memorizes these patterns by synaptic plasticity at the recurrent synapses.

**Fig. 1.**
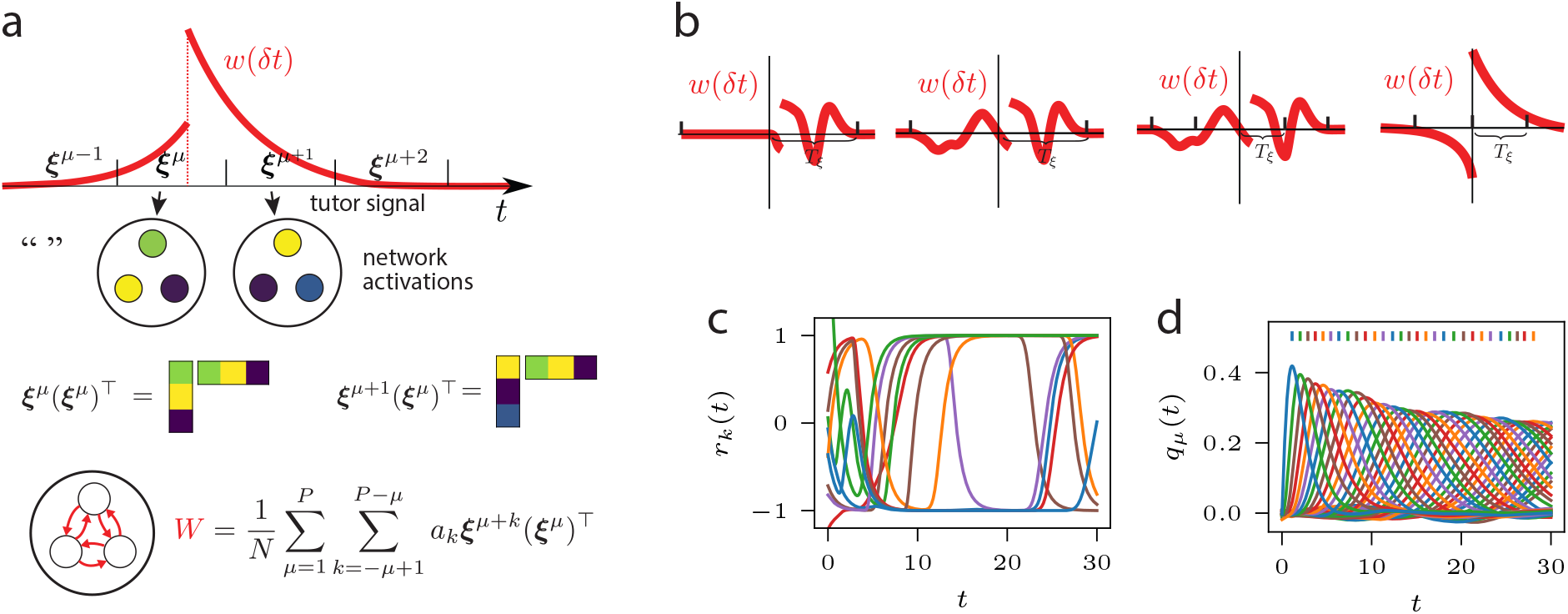
Illustration of network model and stereotypical behavior. **a**) The interaction of a tutor signal (the *ξ*^*μ*^) with Hebbian temporal kernel *w*(*δt*) = *w*(*s* − *t*) gives rise to weight matrix *W*. **b**) Illustrations of different possible kernels. Left: kernel that would give rise to two terms for each 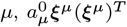 and 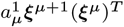. Second from left: kernel that would give rise to three terms for each *μ*, 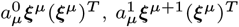 and 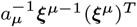. Third from left: kernel that would give rise to five terms for each *μ*, corresponding to *μ* − 2, *μ* − 1, *μ, μ* + 1, and *μ* + 2. Right: double-sided exponential kernel. **c**) Activity *r*_*k*_ of ten randomly selected units in the the network for coefficient values *a*_0_ = 0 and *a*_1_ = 1. **d**) Overlaps *q*_*μ*_ of the network activity with patterns ***ξ***^*μ*^ (here *a*_0_ = 0 and *a*_1_ = 1). Color corresponds to pattern index *μ*. The vertical lines indicate the locations *t*_*μ*_ of the peaks. The first overlap *q*_1_ is not included in the plot for ease of visualization.

To store a memory of this sequence, this synaptic plasticity can take the form of a temporally asymmetric Hebbian (TAH) learning rule activate during application of the tutor signal (31, 32, 37, 44). An example of a TAH learning rule commonly seen in the literature is

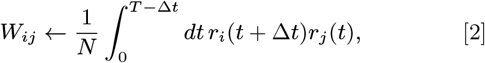

where Δ *t* is an interval of time that spans the amount of time the network is placed in each pattern state and *T* is the duration for the tutor signal associated with the sequence. With this learning rule, connections are strengthened according to the coincident firing of neurons for the current pattern with those of the next pattern; if weights are initially zero, this learning rule results in final weights

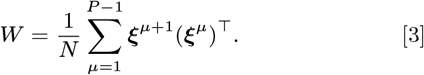

We refer to this weight structure as *sequi*-associative. Intuitively, this set of synaptic weights calculates the overlap between the current network state with the stored patterns, and if the overlap with one of the patterns dominates over others (say pattern *ν*), it steers the network activity to the pattern that is after the dominant pattern (pattern *ν* + 1).

While *sequi*-associative weights capture some temporal aspects of synaptic plasticity, they fall short in their ability to express the full complexity of plasticity dynamics observed in experiments (43, 45, 46) which can significantly affect synaptic structure and network dynamics (47–49). Mathematically, a more general Hebbian learning rule could be written as

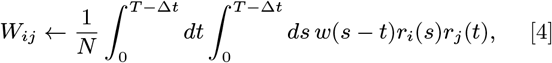

where *w* is a general kernel/filter that weights the Hebbian learning rule as a function of time offset. One sees that Eq. (2) corresponds to *w*(*s* − *t*) = *δ*(*s* − *t* − Δ *t*). However, biologically observed kernels take much richer and different forms, including the famous double-sided exponential STDP curve of Bi and Poo (43) (Fig. 1b, right). Note that the neuroscience literature often plots the kernel as a function of − *δt* = (*s*− *t*) instead of − *δt* = *s*− *t* (i.e. “pre-post” instead of the “post-pre” convention used here).

In this paper, we consider the learning of sequences under a tutor signal with the general TAH rule (Eq. (4)). This rule can give rise to a much richer set of weights defined by

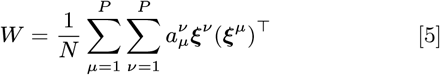

where 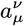 are coefficients that satisfy

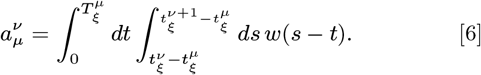

Here 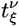 is the time at which the tutor signal for pattern *ν* begins and 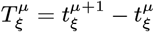 is the duration for which pattern *μ* is shown to the network. Our goal is to understand the sequential dynamics and its tempo arising from such learning rules.

We should note that other works have also considered extensions to Eq. (3). For example, some works (32, 34, 50, 51) include an autoassociative component ***ξ***^*μ*^(***ξ***^*μ*^)^⊤^ to the weights; (32, 34) use discrete dynamics that selectively low-pass filter the *sequi*-associative component, and (50) multiplies the *sequi*-associative term with random noise. The autoassociative term modulates the pattern transition speed. The authors of (40) conduct simulation studies of a model with an additional *prae*-associative term ***ξ***^*μ*− 1^(***ξ***^*μ*^)^⊤^ that has the same strength and opposite sign as the *sequi*-associative component. (35) derive a discrete dynamics in terms of the parameters of an exponentially decaying kernel, giving rise to terms of the form *ξ*^*μ*+*k*^*ξ*^*μ*^ for *k* ≥ 0; however, mathematical analysis of this model is limited to deriving the mean-field dynamics and computing some aspects of stability (there are no calculations related to tempo). (36) similarly consider terms of the form *ξ*^*μ*+*k*^*ξ*^*μ*^, but again focus on capacity and stability properties. Going back to our model, to quantify the sequential behavior of the network dynamics, we define the overlaps

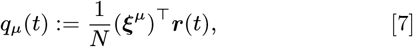

which measure the similarity of the recurrent network activity ***r*** at time *t* to stored patterns ***ξ***^*μ*^. For our initial condition, we will always initialize the network state to the first pattern ***ξ***^1^. Note that the dynamics of the network in the basis of the neurons may not reveal the sequential structure (Fig. 1c), while the overlaps *q*_*μ*_ clearly display this structure (Fig. 1d). To measure sequence progression, we track the location in time *t*_*μ*_ of the maxima of the overlaps *q*_*μ*_(*t*); at time *t*_*μ*_ we say the sequence is in state *μ* (see vertical lines in Fig. 1d).^*^ We also define *p*_*μ*_ := *q*_*μ*_(*t*_*μ*_) to be the height of the peaks. We are particularly interested in the speed of the sequence progression, which we measure by the peak difference *d*_*μ*_ := *t*_*μ*_− *t*_*μ −*1_. Note that the coefficients 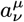 can inhomogeneously depend on *μ*, resulting in a tempo of sequence recall that can be controlled based on the pattern state (i.e. the same sequence can have both fast and slow transitions between patterns, resulting in a *d*_*μ*_ that depends strongly on *μ*). If the tutor signal presents patterns at regular intervals, then the coefficients take the form 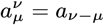. In this case the coefficient for the *ξ*^*ν*^ *ξ*^*μ*^ term only depends on the difference between *ν* and *μ* rather than depending on *μ* independent of *ν*:

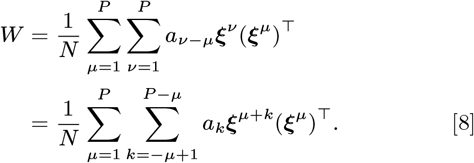

This means that the tempo can no longer be controlled in a sequence-state-dependent (i.e. *μ*-dependent) manner. Even stronger, often the tempo is approximately independent of *μ* (it is a global property of the sequence), so that *d*_*μ*_ is approximately constant across *μ*.

### Sequence stability

Beside tempo, we are also interested in the stability properties of sequential activity. Note that since we are using a saturating nonlinearity *ϕ*, each ***r***_*i*_ is bounded which implies that the overlap *q*_*μ*_ is also bounded. Hence we are primarily interested in whether or not the solutions *q*_*μ*_(*t*) decay to 0 with increasing *μ*, and we say that the sequence of solutions (*q*_*μ*_(*t*))_*μ*_ is stable if the solutions *q*_*μ*_(*t*) do not decay to the zero solution *q*(*t*) = 0 with increasing *μ*.^†^ Formally, in the *T*→ ∞ limit, we say that the sequence is stable if there exists a positive constant *ϵ* such that for all positive integers *μ* there exists a *s*_*μ*_ such that *q*_*μ*_(*s*_*μ*_) > *ϵ*. Note that this definition requires that we take *μ* (and hence *T*) arbitrarily large. If *μ* were bounded in the definition of stability then almost every solution would be stable trivially.

### Mean-field theory

To gain analytical insight about the behavior of our model, we extend the mean-field analysis of (37) to the general weights described by Eq. (5). We assume that the patterns are drawn identically and independently from a Gaussian distribution, 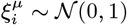, and take the network population size to infinity, *N*→ ∞.

In this mean-field limit the overlaps evolve with dynamics given by (see Appendix 1)

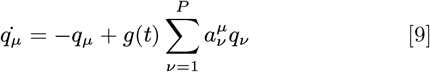

where

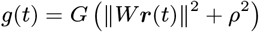

for some function *G*. We give the full functional form of *G*(*x*) in Appendix 1 (Eq. (20)), which depends on the shape of *ϕ*. For the analysis here, the important aspect of *G* is the value of *G*(*ρ*^2^), which serves as an upper bound on *g*(*t*). In particular, for *ρ* = 0, *g*(*t*) is bounded from above by *G*(0), and 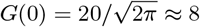 for our choice of parameters for *ϕ*.

The convergence with increasing *N* of the mean-field solutions to the full network solutions is illustrated in Fig. 2. This holds both in the case of uniform tutor signal intervals (Figs. 2a and 2b) as well as non-uniform tutor intervals (Figs. 2c and 2d). For the following, we assume that tutor signal intervals are uniform in temporal duration, so 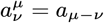. We will revisit the case of non-uniform intervals in the final subsection of Results.

**Fig. 2.**
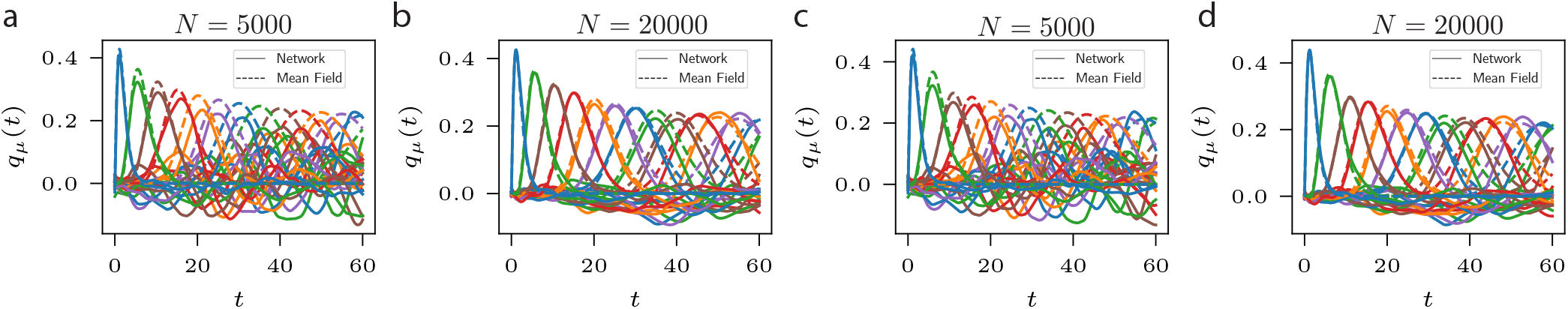
Comparison of full network simulations Eq. (1) (solid lines) with the mean-field equations Eq. (9) (dashed lines). Throughout, every fifth overlap *q*_*μ*_(*t*) is shown, starting with *μ* = 2. **a**) Overlaps *q*_*μ*_(*t*) for network with *N* = 5000 neurons. The only nonzero coefficient is *a*_1_ = 1.5. **b**) As in **a**, but for *N* = 20000 neurons. **c**) As in **a**, but the coefficients are 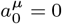 and 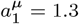 for *μ* < 11; and 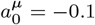 and 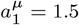 for *μ ≥* 11. **d**) As in **c** but for *N* = 20000.

We next analyze the mean-field theory defined by Eq. (9) to gain insight into the stability properties of sequence generation. To analyze Eq. (9), we use the assumption that *g*(*t*) asymptotes to a constant value over time, and define *g*_∞_ = lim_*t*→∞_ *g*(*t*). We validate this assumption in simulations below.

A fundamental quantity in determining sequence dynamics is 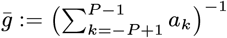 where the sum is over all coefficients. For the following we consider the case of large *N*, and reiterate that these analyses treat the case of uniform tutor signal intervals. In Appendix 2 we show that in this case, and under other mild assumptions, the solutions to the network dynamics Eq. (1) are stable if and only if 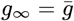. Further, we reason and observe that the network will endeavor to satisfy this stability condition, so that 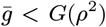 is necessary and sufficient for 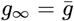. Motivated by this, we define 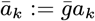 and consider the linear equations

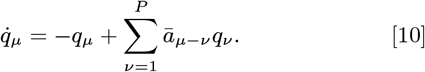

Eq. (10) can be expected to approximate Eq. (9) when the solutions to Eq. (9) are stable (equivalently, when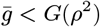). When the solutions to Eq. (9) are unstable (when 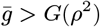), these solutions will vanish with increasing *μ* while the solutions to Eq. (10) will remain stable. We will test the goodness of the linear approximation Eq. (10) to Eq. (9) in simulations.

Note that 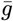 can be related to the TAH kernel *w* in a simple way. To see this, we first use Eq. (6):

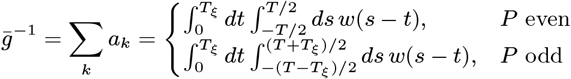

Assuming that *w*(*δt*) becomes vanishingly small for large |*δt*| and that *T* ≫*T*_*ξ*_, the inner integral can be approximately replaced with one over the real line:

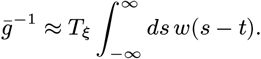

Note that this equation means the network can be stabilized by extending the duration of the tutor signal *T*_*ξ*_. Therefore, a biologically-plausible solution to embedding stable sequences is by extending the tutor signal duration. As we will show below, increasing 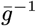 also increases the robustness of the sequence dynamics to noise.

### One forward term

We next turn to analyzing sequence progression in the recurrent neural network. We start by considering the case with one forward term of Eq. (10), where *a*_*k*_ = 0 for *k* ∉ {0, 1}. This case is relevant in the regime where the duration of sequence elements, *T*_*ξ*_, is significantly longer than the timescale of *w*(*t*) (i.e. *w*(*δt*) ≈0 for *δt* > *T*_*ξ*_ and *w*(*δt*) = 0 for *δt* < 0). Figure 3a illustrates two possible kernels *w* corresponding to this case.

**Fig. 3.**
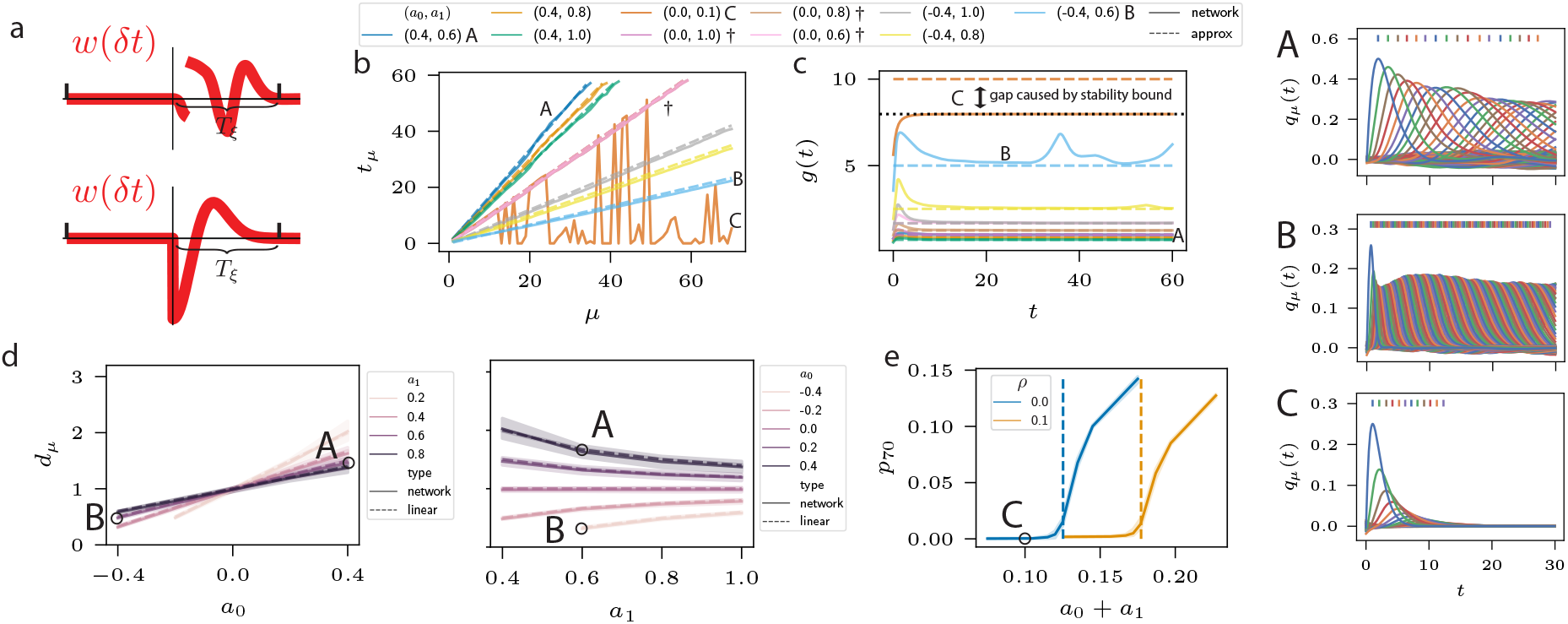
Sequence progression for two-term connectivity *W*^*μ*^. Plots show the full network simulations (solid lines) and linear approximations Eq. (11) (dashed lines). Throughout, capital letters *A, B*, and *C* correspond to parameter values as indicated in the legend. **a**): Illustrations of two TAH kernels *w* that would give rise to two terms. **b**) Peak times *t*_*μ*_. The *†* symbol denotes parameter values that give rise to overlapping lines, and C denotes a parameter value for which the network equations are unstable. **c**) The value of *g*(*t*) through time during the simulations. Dashed lines denote 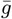. The function *g*(*t*) is bounded from above by *G*_max_ *≈* 8, which is indicated by a dotted line. For the parameters corresponding to *C* there is a mismatch between 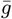 and *g*_∞_ (asymptote of solid line). **d**) Plots of peak differences *d*_*μ*_ as a function of *a*_0_ and *a*_1_. Shaded regions are 95% confidence intervals for *μ* ∈ {2, …, 10}, and lines are the means. Left: *a*_0_ is plotted on the horizontal axis and *a*_1_ is denoted by color. Right: *a*_1_ is plotted on the horizontal axis and *a*_0_ is denoted by color. **e**) The peak height *p*_70_ corresponding to the maximum of *q*_70_(*t*) (*μ* = 70) over *t*, in the full network simulation, as a function of the sum of coefficients *a*_0_ + *a*_1_. Color denotes the noise strength *ρ* and dashed vertical lines show the critical points *G*(*ρ*^2^)^− 1^ where the network is predicted to pass from stable to unstable. Shaded regions are confidence intervals over *a*_0_ ∈ {− 0.2, 0, 0.1}. **A**) Overlaps *q*_*μ*_(*t*) corresponding to parameter values (0.4, 0.6). **B**) Overlaps *q*_*μ*_(*t*) corresponding to parameter values (− 0.4, 0.6). **C**) Overlaps *q*_*μ*_(*t*) corresponding to parameter values (0.0, 0.1).

In this case Eq. (10) becomes

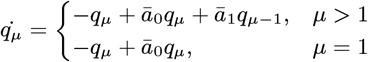

The solutions to these equations are (recall the initial conditions *q*_*μ*_(0) = *δ*(*μ* − 1))

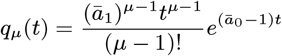

which have maxima at

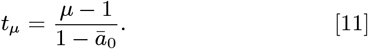

Note that *ā*_0_ + *ā*_1_ = 1 by definition, which places some restrictions on the values that *ā*_0_ and *ā*_1_ can take. For instance, *ā*_0_ > 1 implies *ā*_1_ < 1. This ensures that 0≤ *q*_*μ*_(*t*) ≤1 for all choices of *a*_0_ and *a*_1_. The (inverse) speed as measured by peak difference is

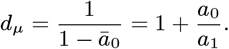

Figure 3 shows the simulation results for various values of *a*_0_, *a*_1_, and *ρ*. Figure 3b plots peak times *t*_*μ*_ for *ρ* = 0. The full network simulations with *N* = 30000 match the linear approximations Eq. (11) closely, except for parameter values where the solutions for the full network decay to zero (denoted by C). This happens because 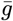 is larger than the critical value of *G*(0) ≈8. Note the significant role of *a*_0_ in determining the sequence speed. In particular, for *a*_0_ = 0 changing *a*_1_ has no impact on the speed († symbol in Fig. 3b). Note also that, unlike in standard Hopfield models, the autoassociative term *a*_0_ can be negative. The bottom panel of Fig. 3a shows an example of a kernel that gives rise to negative *a*_0_ and positive *a*_1_.

Figure 3c plots the function *g*(*t*). This plot supports our original assumption that *g*(*t*) asymptotes to a constant value, and shows the close match between *g*_∞_ and 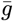, except in the case where 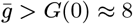.

To look more closely at the relative impacts of *a*_0_ and *a*_1_ on sequence speed, we plot sequence speed for different combinations of *a*_0_ and *a*_1_ in Fig. 3d. Figure 3d shows that the dependence of *d*_*μ*_ on *a*_0_ is roughly linear in the range plotted, with a slope that decreases with *a*_1_. Parameter ranges were chosen in an attempt to capture the full behavior of the dynamics within the region of stable sequence generation; in particular, this requires that *a*_1_ > 0 and *a*_0_ + *a*_1_ > 1*/G*(0). Note that for *a*_0_ < 0 sequence speed decreases, counter-intuitively, with increasing *a*_1_, while the opposite relationship holds for *a*_0_ > 0. This can be seen from the expression for *d*_*μ*_ above.

Figure 3e quantifies the stability properties of sequences by plotting the peak magnitude *p*_*μ*_ for a pattern of large index (here *μ* = 70) as a function of 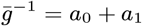. Color denotes the strength of noise injected into the network dynamics (*ρ* in Eq. (1)). Note that such noise could come from interference, such as that which arises if there are multiple stored sequences (37). The dashed vertical lines mark the critical values of stability where 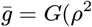). This plot illustrates how 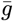 determines the stability of sequences, and that smaller 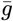 (equiv. larger *a*_0_ + *a*_1_) is required for stability in the presence of noise. The plot shows that, as predicted, peak magnitudes *p*_*μ*_ have a sharp inflection point at 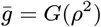, where *p*_*μ*_ goes from being approximately zero to nonzero.

Note that *a*_0_ and *a*_1_ can be adjusted so that sequence progression speed *d*_*μ*_ remains constant, but stability is improved by decreasing 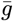. This can most easily be seen by considering lines of constant *d*_*μ*_ in Fig. 3d. The most straightforward example is the line *d*_*μ*_ = 1 in the right panel of Figure 3d, which clearly corresponds to many values of *a*_0_ + *a*_1_. This is particularly important in the presence of noise, as shown by Fig. 3e. This degree of freedom of the system makes some kernels *w*(*t*) strictly better than others, even if they result in the same speed.

### General approximate solution

While we were able to explicitly solve Eq. (10) in the case of two nonzero terms in the preceding section, in general an explicit solution is not available and we need to resort to approximations. To approximately solve Eq. (10), we replace the equation with one that is periodic in *μ*. A general approximate solution to the equations with these periodic boundary conditions can then be found (see Appendix 2):

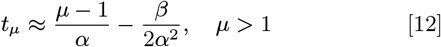

where 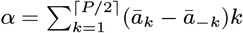 and 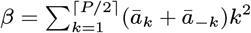. Sequence speed as measured by peak difference is then

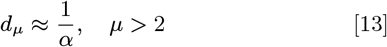

When *α* > 0, these equations to describe forward propagation of the sequence.

In the following sections we will look at special cases of interest that illustrate the theory.

### Three terms with bidirectional connectivity

Next we consider the case with three terms and bidirectional connectivity in Eq. (10), where *a*_*k*_ = 0 for *k* ∉ {− 1, 0, 1}. Bidirectional connectivity occurs with TAH kernels commonly found in biology, such as the double-sided exponential decay kernels (52) (see Fig. 4a for an illustration). Three terms in particular are relevant when the TAH kernel’s timescale is faster than the tutor signal’s, for instance when the support of the kernel vanishes outside of time windows spanning more than two pattern presentations. Figure 4a illustrates a temporally asymmetric kernel that would give rise to such three terms.

**Fig. 4.**
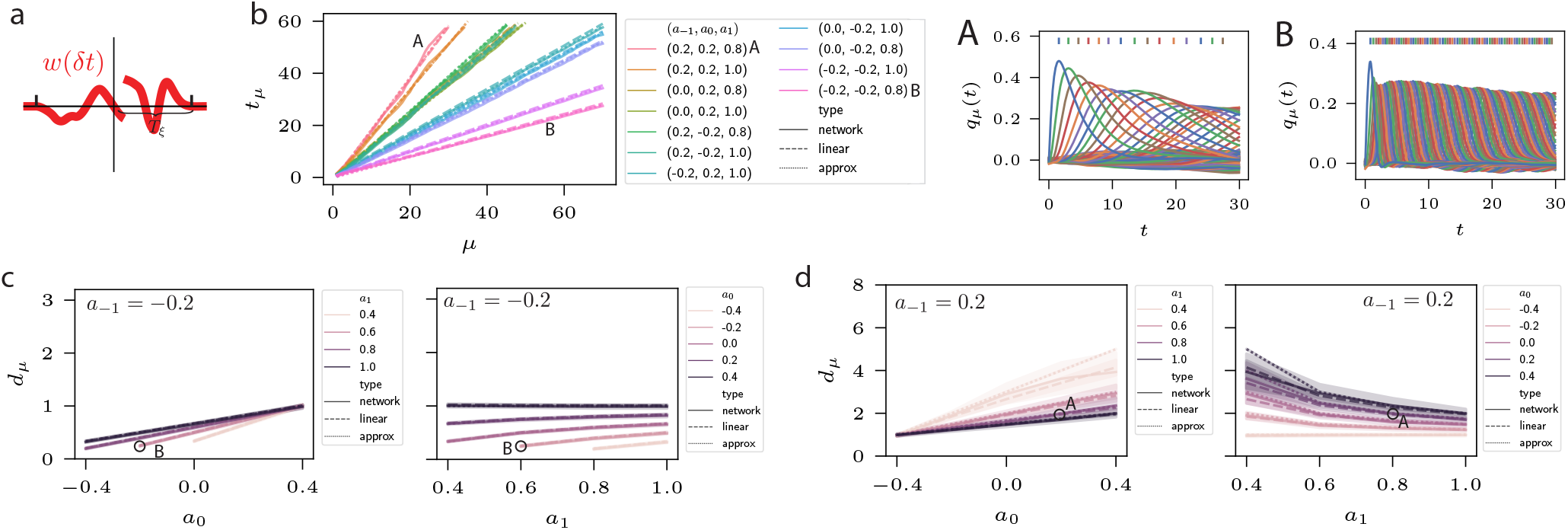
Sequence progression for bidirectional terms. Plots show the full network simulations (solid lines), linear approximations Eq. (10) (dashed lines), and approximation Eq. (14). Throughout, capital letters *A* and *B* correspond to parameter values as indicated in the legend. **a**) Illustration of a TAH kernel *w* that would give rise to bidirectional terms. **b**) Peak times *t*_*μ*_ plotted for a variety of coefficient combinations (*a*_− 1_, *a*_0_, *a*_1_), denoted by color. **c**) Plots of peak differences *d*_*μ*_ as a function of *a*_0_ and *a*_1_. Here *a*_− 1_ = *−* 0.2. Shaded regions are 95% confidence intervals for *μ* ∈ {2, …, 10}, and lines are the means. Left: *a*_0_ is plotted on the horizontal axis and *a*_1_ is denoted by color. Right: *a*_1_ is plotted on the horizontal axis and *a*_0_ is denoted by color. **d**) As in **c** but with *a*_− 1_ = 0.2. **A**) Overlaps *q*_*μ*_(*t*) corresponding to parameter value (0.2, 0.2, 0.8). **B**) Overlaps *q*_*μ*_(*t*) corresponding to parameter value (− 0.2, − 0.2, 0.8).

To understand sequence progression speed, we use Eq. (12):

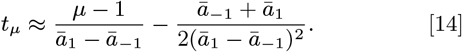

Sequence speed as measured by peak difference is then

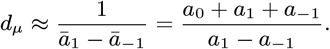

The effect of changing *a*_− 1_ on sequence speed is revealed by taking the derivative of *d*_*μ*_ with respect to *a*_− 1_:

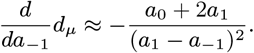

This shows that increasing *a*_− 1_ decreases *d*_*μ*_ provided *a*_0_ + 2*a*_1_ > 0. To compare the relative impact of changing *a*_− 1_ and *a*_1_ on sequence speed, we can take the ratio of derivatives:

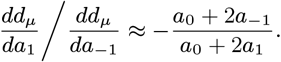

This equation reveals that changes in *a*_− 1_ will impact sequence speed more than changes in *a*_1_ when *a*_− 1_ is smaller in magnitude than *a*_1_. Hence *a*_− 1_ is a natural parameter to use to control speed.

Figure 4 compares the full network simulations, the simulations of the linear system Eq. (10), and the approximation Eq. (14). The peak times *t*_*μ*_ are plotted in Fig. 4b, showing that Eq. (14) is a good estimate of both the linear and non-linear systems. Figure 4c shows the relationship between *d*_*μ*_, *a*_0_, and *a*_1_ for fixed *a*_− 1_ = − 0.2. The left panel reveals that *d*_*μ*_ increases approximately linearly with *a*_0_, with a slope that decreases with *a*_1_. Comparison with Fig. 3d shows the impact of introducing *a*_− 1_, which can be mixed but is generally to decrease *d*_*μ*_ over the parameter regime considered.

Figure 4d shows the relationship between *d*_*μ*_, *a*_0_, and *a*_1_ for fixed positive *a*_− 1_ = 0.2. Again, the left panel reveals that *d*_*μ*_ increases approximately linearly with *a*_0_, with a slope that decreases with *a*_1_. Comparing with Fig. 3d shows how *a*_− 1_ = 0.2 typically increases *d*_*μ*_ relative to *a*_− 1_ = 0.

As in the case with two terms, the speed of the sequence can be held constant while the stability term 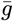 is decreased. Indeed, the introduction of *a*_− 1_ introduces an additional degree of freedom. Examples of keeping *d*_*μ*_ fixed while varying 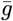 can be seen by following horizontal lines of fixed *d*_*μ*_ in Figs. 4c and 4d.

### Three terms with forward connectivity

Now we consider the case with two forward terms, where *a*_*k*_ = 0 for *k* ∉ 0, {1, 2}. This illustrates the effects of having slower timescale TAH kernels, roughly equal to or a bit slower than the timescale of the tutor signal. Figure 5a illustrates a TAH kernel that would give rise to two forward terms. Using Eq. (12):

**Fig. 5.**
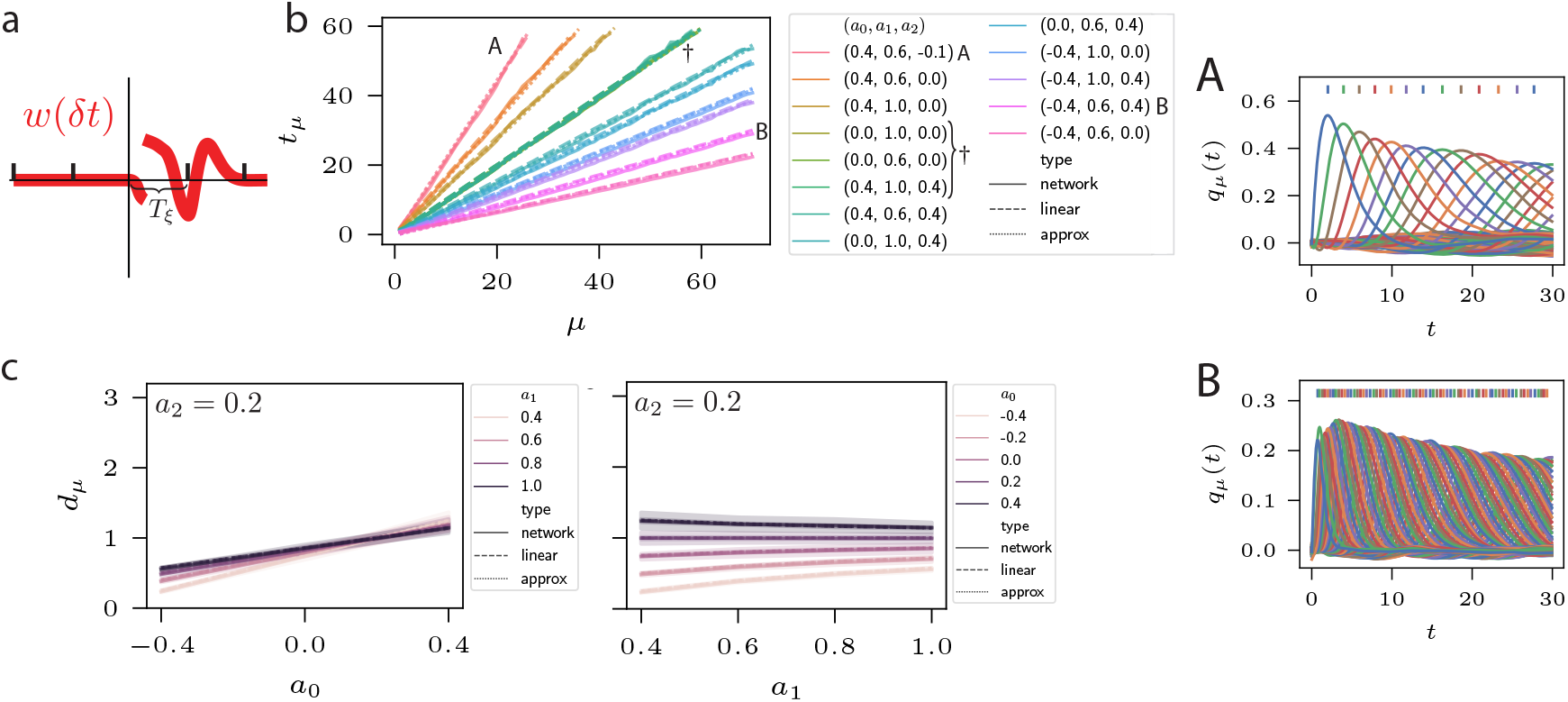
Sequence progression for two forward terms. **a**): illustration of a TAH kernel *w* that would give rise to two forward terms. Plots show the full network simulations (solid lines), linear approximations Eq. (15) (dashed lines), and approximation Eq. (12) (dotted lines). **b**) Peak times *t*_*μ*_ for a variety of coefficient combinations (*a*_0_, *a*_1_, *a*_2_), denoted by color. Dotted lines are the linear approximation given by Eq. (15). The *†* symbol denotes parameter values that give rise to the same peak times. **c**) Plots of peak differences *d*_*μ*_ as a function of *a*_0_ and *a*_1_. Here *a*_2_ = 0.2. Shaded regions are 95% confidence intervals for *μ* ∈ {2, …, 10}, and lines are the means. Left: *a*_0_ is plotted on the horizontal axis and *a*_1_ is denoted by color. Right: *a*_1_ is plotted on the horizontal axis and *a*_0_ is denoted by color. **A**) Overlaps *q*_*μ*_(*t*) corresponding to parameter value (0.4, 0.6, − 0.1). **B**) Overlaps *q*_*μ*_(*t*) corresponding to parameter value (− 0.4, 0.6, 0.4).

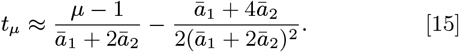

Sequence speed as measured by peak difference is then

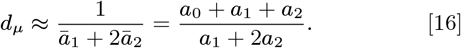

Figure 5 compares the full network simulations, the simulations of the linear system Eq. (10), and the asymptotic approximation of the peak times Eq. (15), showing a close match for all three quantities. Similar to the case of the previous section, increasing *a*_2_ has a qualitatively different effect on *d*_*μ*_ depending on the values of *a*_0_ and *a*_1_. Taking the derivative of *d*_*μ*_ with respect to *a*_2_ in Eq. (16) –

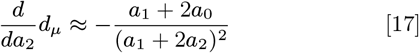

– indicates that increasing *a*_2_ increases *d*_*μ*_ if and only if *a*_1_ + 2*a*_0_ < 0. Unlike in the previous section, this condition cuts through a significant portion of the region of coefficient space where stable, forward propagating sequences are viable. Examples of the effect of increasing *a*_2_ can be seen in Fig. 5b, which plots peak times *t*_*μ*_ for a range of coefficient combinations. The differing effect of increasing *a*_2_ can be seen in this plot.

The relationship between *d*_*μ*_ and different coefficient combinations is further elucidated in Fig. 5c, which plots *d*_*μ*_ as a function of *a*_0_ and *a*_1_ where *a*_2_ = 0.2. As in previous sections, the dependence of *d*_*μ*_ on *a*_0_ is approximately linear over the coefficient values considered, with a slope that decreases with *a*_1_. Comparison with Fig. 3d shows the (multiplexed) effect of positive *a*_2_.

Note that, as in the previous sections, the stability properties of the sequence (determined by the sum of coefficients) can be improved even as the timing is held constant (†symbol in Fig. 5b); that is, 1*/*(*ā*_1_ + 2*ā*_2_) can be fixed while 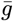 is decreased. This is most clearly illustrated by the horizontal line *d*_*μ*_ = 1 in the right panel of Fig. 5c.

Here we consider the special case of (double-sided) exponential kernels as commonly used to model STDP kernels seen in biology (43, 52). These have the form 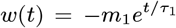 for *t* < 0 and 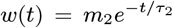 for *t* ≥ 0 (see Fig. 6a for an illustration).

**Fig. 6.**
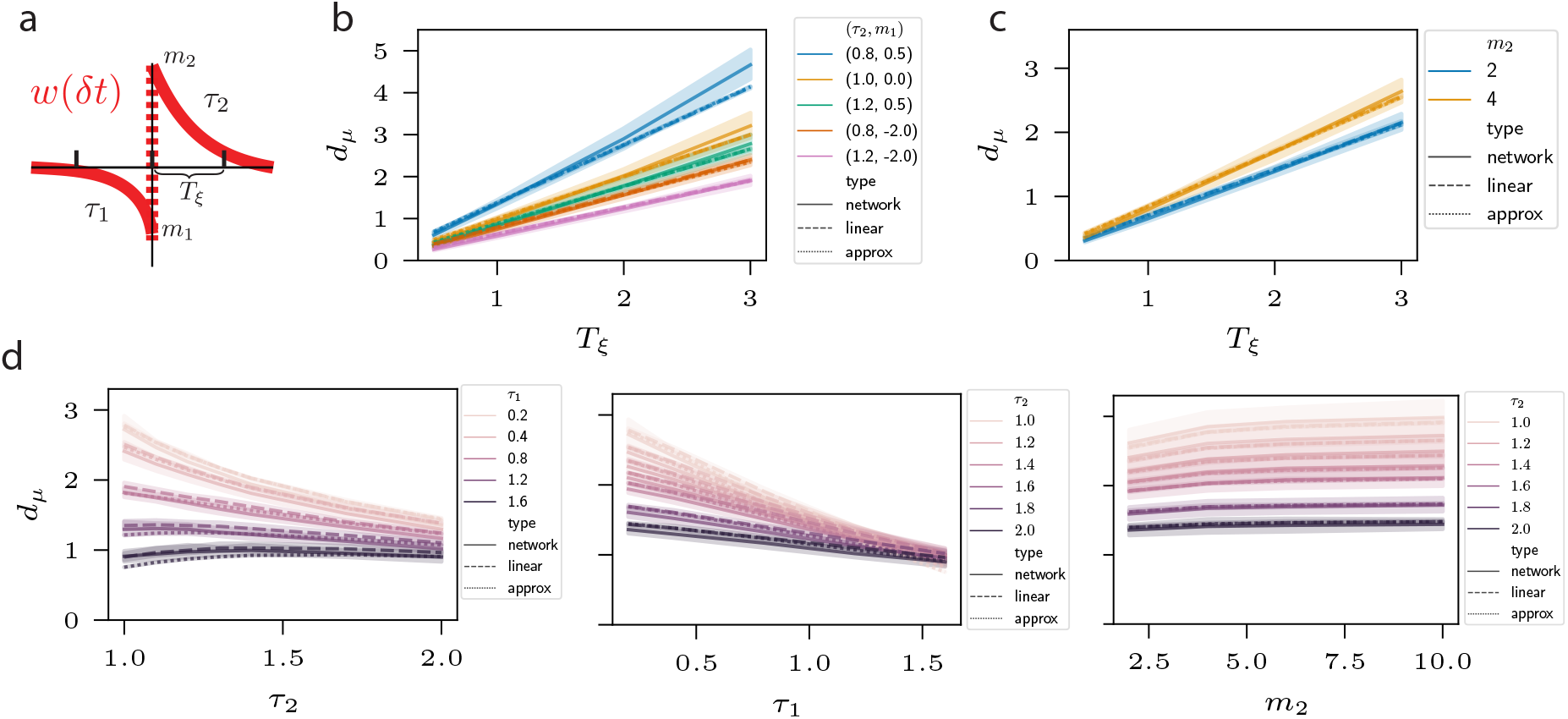
Sequence progression for double-sided exponential kernels and varying *T*_*ξ*_. Shaded regions are 95% confidence intervals over *μ ∈ {*2, …, 10*}*, and lines are the means. Plots show the full network simulations (solid lines), linear approximations Eq. (10) (dashed lines), and approximation Eq. (12) (dotted lines). **a**): Illustration of a double-sided exponential TAH kernel *w*. **b**) Peak differences *d*_*μ*_ = *t*_*μ*_ − *t*_*μ*− 1_ for a variety of double-sided exponential TAH kernel parameters *τ*_2_ and *m*_1_ (color). Here *τ*_1_ = .25 and *m*_2_ = 2. **c**) As in **b**, but with *m*_1_ = − 2 and changing *m*_2_ (denoted by color). **d**) Peak differences *d*_*μ*_ as a function of *τ*_1_, *τ*_2_, and *m*_2_. Unless otherwise defined, parameter values are *τ*_1_ = .25, *m*_1_ = − 1, *m*_2_ = 3, and *T*_*ξ*_ = 3. Left: *d*_*μ*_ as a function of *τ*_2_ with varying *τ*_1_ (color). Center: *d*_*μ*_ as a function of *τ*_1_ with varying *τ*_2_ (color). Right: *d*_*μ*_ as a function of *m*_2_ with varying *τ*_2_ (color).

### Exponential kernels

We compute in Appendix 2 that the approximate peak times are given by Eq. (12) where

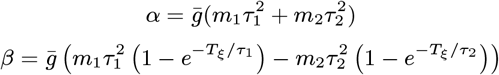

and

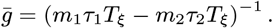

Here *T*_*ξ*_ is the length of the tutor signal interval. Note in particular that the (inverse) sequence speed as measured by

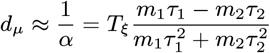

scales linearly with *T*_*ξ*_. This dependence provides a biologically-plausible mechanism to control the speed of recalled sequence by the tutor.

The quantity *d*_*μ*_ is plotted in Figs. 6b and 6c for a variety of different kernels as a function of tutor interval *T*_*ξ*_. These plots compare the full network simulations with the linear approximation Eq. (10) and the approximation Eq. (12), showing a close match, especially for small *T*_*ξ*_.

In Fig. 6d we plot the dependence of inverse sequence speed *d*_*μ*_ as a function of *τ*_1_, *τ*_2_, and *m*_2_. The left panel shows that *d*_*μ*_ typically decreases with increasing *τ*_2_, but this can be reversed for large enough *τ*_1_. The center panel shows that, for the parameter values chosen, *d*_*μ*_ decreases with *τ*_1_. The right panel shows the influence of changing *m*_2_ on *d*_*μ*_; while increasing *m*_2_ increases *d*_*μ*_, the effect is small. Parameter values were chosen to demonstrate the full spectrum of behaviors within the regime of stable, forward propagating dynamics.

To look more closely at the interplay between the timescale of the TAH kernel and the timescale of the tutor signal, we look at the case where *τ*_1_ is small relative to *τ*_2_. Then *α*≈ *τ*_2_*/T*_*ξ*_, indicating that in this case the ratio of the timescales sets the sequence speed.

### Sequences with fast and slow parts

In naturalistic settings, different states of a sequence may have different duration. For instance, in playing a piece of music different notes are held with different durations. Our analysis indicates that changing these durations causes a change in the sequence progression speed, meaning that variable duration tutor signals 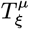 will result in sequences with faster and slower parts (Fig. 7). Consider a sequence with sections that pass from one set of uniform durations 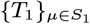 to another 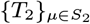. The overlaps *q*_*μ*_ for *μ* ∈*S*_1_ will be governed by the first interval length, with some disturbance from *q*_*ν*_ for *ν* ∈*S*_2_ mediated by the backward coefficients *a*_*k*_ for *k* > 0. For *μ* far from the boundary this disturbance should be small. On the other hand, *q*_*ν*_ far from the boundary will be governed by the second interval length with small disturbance from *q*_*μ*_, with the caveat that the evolution up to time *t* has been influenced significantly by the dynamics of *q*_*μ*_. With these caveats in mind, we should expect the speed in each of these regions far from the boundary to progress at speeds determined by the corresponding tutor signal interval length, with more complex behavior occuring near the boundary of the transition. In Fig. 7 we consider a case with two transitions, where *T*_1_ = 1 for *μ*∈ [1, 20], *T*_2_ = .5 for *μ*∈ [21, 30], and *T*_3_ = 1 for *μ*∈ [31, 80] (slow to fast back to slow). Here we use a double-sided exponential kernel with *τ*_1_ = 0.5, *m*_1_ = 5, *τ*_2_ = 1, and *m*_2_ = 8. This figure illustrates that after transitioning, the sequence speeds up or slows down to match the new duration.

**Fig. 7.**
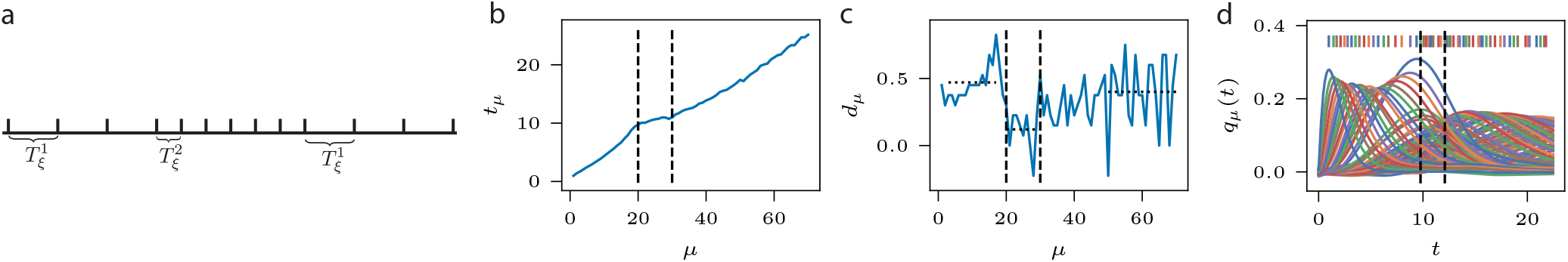
Sequences with fast and slow parts. Sequence dynamics with three regions 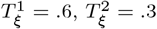, and 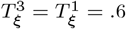. The first transition occurs at *μ* = 20 and the second at *μ* = 30. Parameters are *τ*_1_ = 0.5, *m*_1_ = − 2, *τ*_2_ = 1, and *m*_2_ = 2. **a**) Illustration of the three regions (note that this schematic is not accurate with respect to the *μ* values at which transitions occur). **b**) Peak times as a function of pattern index *μ*. Dashed vertical lines demarcate the three regions. **c**) Peak difference as a function of pattern index *μ*. Dotted lines mark averages over the intervals [3, 18], [20, 28], and [50, 80]. **d**) Plot of overlaps *q*_*μ*_(*t*). Dashed vertical lines demarcate the three regions.

However, these transitions have undesirable characteristics. For one, in the first transition the sequence slows down just before entering the faster sequence section. A second issue is that the peak differences become more variable after each transition. A third issue is that the transition from 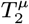 to 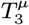 is very gradual and takes a significant amount of time before recovering the speed of the 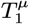 section. These issues, and particularly the third, are probably related to how the overlaps *q*_*μ*_ “spread out” for increasing index *μ* (see Fig. 1 for an example); that is, the width of the bumps traced out by *q*_*μ*_(*t*) increases. This suggests a modified sequence model where the behavior of the model is agnostic to the index of the pattern being represented, perhaps with the use of nonlinear learning rules as in (37, 51) or “interaction modulation” as suggested by (53).

## Discussion

The ability to store experiences as memories is fundamental to intelligent behavior. While memories are typically modeled as fixed points in recurrent networks, sequences constitute information that is also of primary importance. This is clear when considering the ubiquity of well-practiced, automatic motor behaviors.

Our model stores memories by assuming that the network is placed in states by a “tutor signal” during the learning process. There are multiple possible interpretations for this. The state of the network could be set by sensory inputs, so that TAH learning rules constitute an unsupervised method for learning sequential structure in the world. Another interpretation is that higher-level brain areas provide this tutor signal. For instance, motor cortex could be directing motor outputs (a process involving slow, deliberate thought) while sending tutor signals to a subcortical network which makes a copy of the behavioral control that allows sequential behavior to be later recalled, thereby skipping the slow deliberation process and corresponding to automatic behavior (54, 55).

This work seeks to fill a significant gap in theory describing the ability to form sequence memories. Namely, it is not clear how general and commonly observed Hebbian learning rules could give rise to such sequence memories. Hebbian learning rules have the advantage of being simple, biologically plausible, implementable in neural circuits, and continuously differentiable (as opposed to, say, storing data in a database). Our theory may also be useful for analysis of dynamical systems with sequence dynamics more generally, since the mean-field equations Eq. (9) are quite general.

Our analysis reveals many interesting phenomena. For one, Hebbian kernels can be chosen that produce the same sequence (in terms of timing), but that are more or less robust to noise. Our theory lays out a quantitative framework that allows such kernels to be delineated. We derive simple relationships between the parameters of the (double-sided) exponential kernels commonly seen in biology, and show that sequence timing scales approximately linearly with the timing of the tutor signal, and the slope of this linear relationship depends on the parameters of the kernel. Finally, we show the result of changing the timing of the tutor signal throughout the sequence, so that the sequence has faster and slower parts. While the sequence timing roughly follows these changes, we find several characteristics of these changes that leave something to be desired and suggest possible goals for improving the model.

A shortcoming of our model is that the possibility of correlated patterns is not addressed. It is reasonable to think that sequence state patterns could be correlated in biological circuits, and addressing this may provide a more useful and biologically faithful model. An additional limitation is the interpretation of discrete sequence states, and on understanding what the biologically relevant timescales for tutor signals are. It is possible that neural control signals in higher order brain areas are “chunked” – that is, a relatively constant control signal initiates another motor area that enacts a motor motif (1). In addition, we consider only linear TAH learning rules, although we expect our analysis would extend in spirit to nonlinear learning rules such as those considered in (37).

In the longterm, memory models should be combined with other models, such as models of motor cortex supporting flexible motor control. Tutor signals would then be generated by these other models, and gradually the memory model would take over control of motor actions as they become more practiced (56). In addition, mechanisms for refinement based on reward signals could be added to the memory modules. For instance, by gating TAH learning rules based on reward signals, this learning process can be turned into a reinforcement learning process.

In summary, our analysis shows that sequence memories can be formed by simple biologically plausible learning rules with general TAH kernels and sequence speed is set by the interplay between the TAH kernels and the duration that the network is tutored. Our work indicates areas of potential improvement and future development that we believe are important for advancing the field.

## Acknowledgements

MF thanks the Swartz Foundation and HDSI for its support. CP is supported by NSF grant DMS-2134157.

## Code availability

Code for reproducing the plots can be found at https://github.com/msf235/Recall-tempo-of-Hebbian-sequences. For details of the parameters used in the simulations for producing the plots, see Appendix 3.

## Appendix 1

### Derivation of mean-field equations

Here we derive the mean-field equations Eq. (9). This is a fairly straightforward generalization of the derivation in (37). The full network equations are 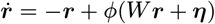 where ***η***(*t*) is a white noise vector (⟨ *η*_*i*_(*t*)⟩ = 0 and ⟨ *η*_*i*_(*t*)*η*_*i*_(*t*^′^)⟩ = *ρ*^2^*δ*(*t*− *t*^′^)) and *ϕ* is a sigmoidal nonlinearity:

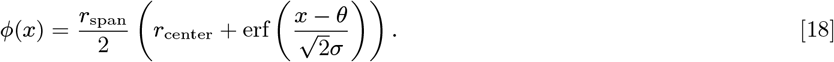

The weights *W* are

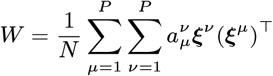

Here each pattern ***ξ***^*μ*^ is standard normal: 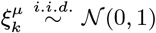. The mean-field equations are written in terms of the overlaps *q*_*μ*_(*t*) = (***ξ***^*μ*^)^⊤^ ***r***(*t*)*/N*. If we let ***h*** = *W* ***r*** + ***η*** we can then write

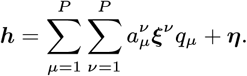

We now investigate the evolution of the overlaps *q*_*μ*_:

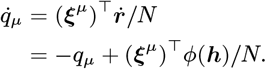

Let’s look more closely at the second term on the right hand side. As *N* → ∞ this term approaches an average by the law of large numbers, yielding (***ξ***^*μ*^)^⊤^ *ϕ*(***h***)*/N* → ⟨ *ξ*^*μ*^*ϕ*(*h*)⟩ where 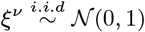 and

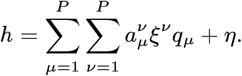

Here *η* is a scalar white noise term: ⟨ *η*(*t*)⟩ = 0 and ⟨ *η*(*t*)*η*(*t*^′^)⟩ = *ρ*^2^*δ*(*t* − *t*^′^). Hence,

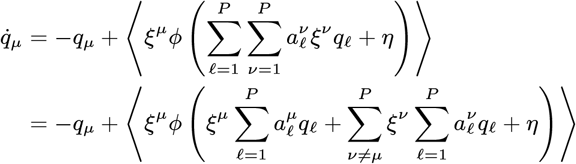

We next note that each term of

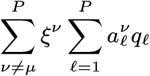

is an independent normally distributed random variable with mean zero and variance 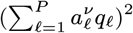. Hence the sum define 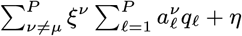 is distributed like *xR*_*μ*_ where 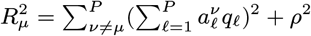 and *x* is standard normal. We similarly 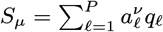 and write the average as a Gaussian integral:

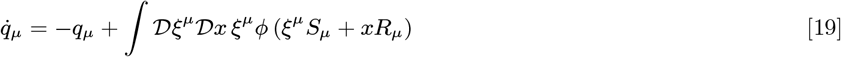

where 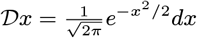.

The remainder of the calculation follows closely (37). We next define the change of coordinates

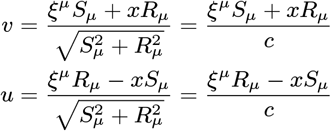

where *v* and *u* are uncorrelated standard normal random variables and where we have defined 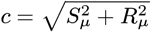. To perform this change of variables we first compute the determinant of the Jacobian:

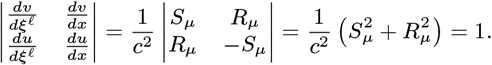

Hence we can use the substitution *dudv* = *dξ*^*μ*^*dx*. Furthermore,

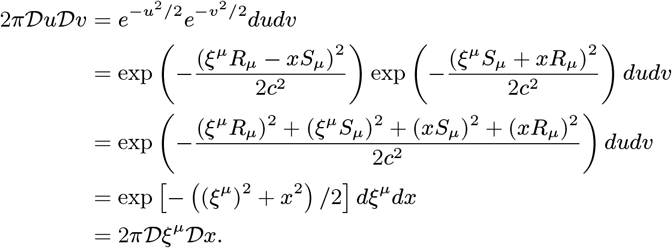

Inverting the change of coordinate equations above, we find that *ξ*^*μ*^ = (*S*_*μ*_*v* + *R*_*μ*_*u*)*/c*. Using these substitutions, we can rewrite the integral

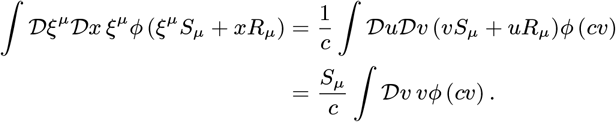

We now define

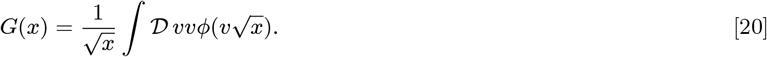

Using the definition of *ϕ*, this integral can be evaluated with integration by parts and is

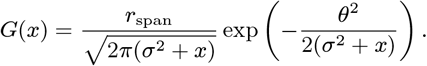

Our equation becomes

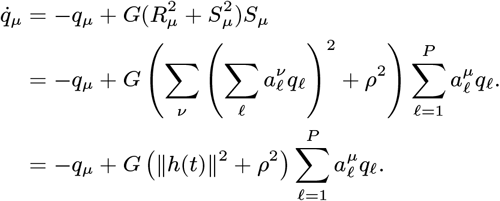

Finally, we define

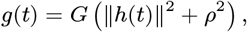

which results in Eq. (9). Plots showing the convergence of the mean-field approximation to the network simulations for increasing *N* are shown in Fig. 2. Note that in subsequent analysis, the number of patterns *P* will also be taken to infinity. In order for these mean-field dynamics to hold exactly, *P/N* must vanish as *N* and *P* are both taken large.

Note that stability is determined by the shape of the function *G*. For the values of *ϕ* considered in the main text (*r*_span_ = 2, *r*_center_ = 0, *θ* = 0, and *σ* = 0.1), *G* has the shape plotted in Fig. S1. In particular, *G* is monotonically decreasing from its maximum at *x* = 0. Since ∥*h*(*t*) ∥^2^ ≥ 0, the maximal value that *g*(*t*) can take during the ongoing dynamics is bounded from above by *G*(*ρ*^2^). Note that the function *G* is the same as in (37), and the supplementary material of (37) contains plots of *G* for various values of *r*_span_, *θ*, and *σ*.

## Appendix 2

In this appendix we show the full details of the mathematical derivations of the peak times in the main text.

**Fig. S1.**
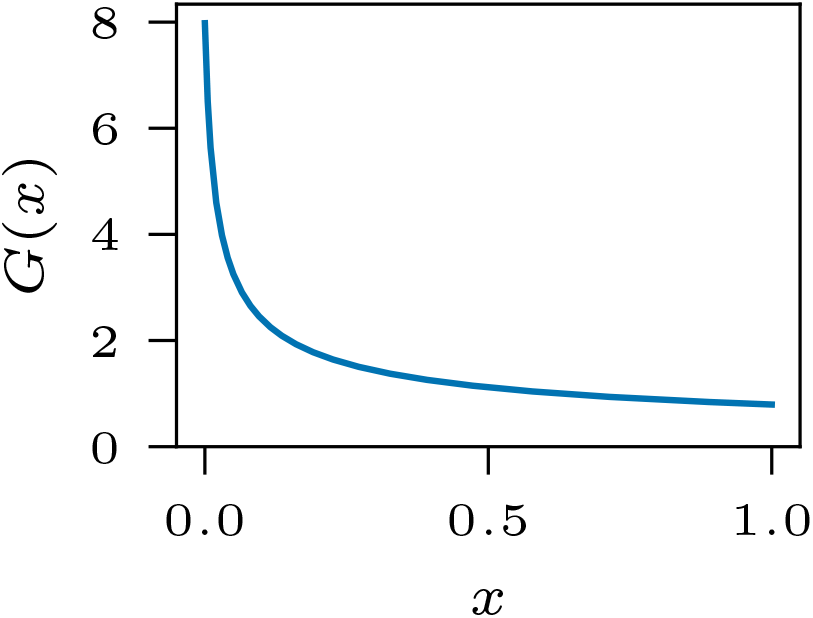
The function *G*(*x*) for *r*_span_ = 2, *θ* = 0, and *σ* = 0.1.

### Derivation of *g*_∞_

Here we derive how the value of *g*_∞_ = lim_*t*→∞_ *g*(*t*) depends on the coefficients *a*_*k*_ and the nonlinearity *ϕ*. We start with the mean-field equations Eq. (9) and assume that the coefficients are shift-invariant, so that 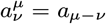s:

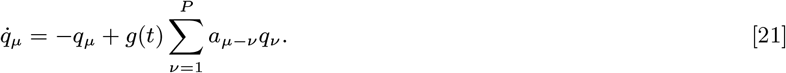

We show that

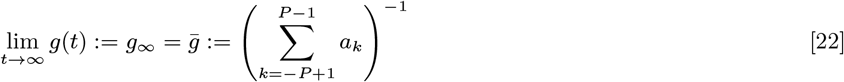

provided the solution to Eq. (21) is stable, and under a few other mild assumptions. First, we assume that *g*_∞_ exists. Using this assumption, we note that for large enough *t* the equations Eq. (21) become approximately linear (since *g*(*t*) becomes approximately constant). Hence we can approximate the limiting behavior of Eq. (21) by the linear system

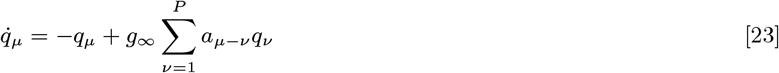

provided we use as initial condition for Eq. (23) the solution of Eq. (21) evaluated at some large enough *t*. Since the limiting behavior (in terms of stability) of the linear system Eq. (23) does not depend on initial condition, this choice of initial condition is not of substance. Hence to analyze the asymptotic stability properties of Eq. (21), we can instead analyze the asymptotic stability properties of Eq. (23)

To analyze the asymptotic stability properties of Eq. (23), we instead consider solutions to this equation with periodic boundary conditions (in *μ*). While this change in boundary conditions may influence the final solution, it should not affect the asymptotic stability behavior as we take *P* → ∞. This change in boundary conditions requires approximating our kernel function *w*(*t*) with a kernel function that is *T*-periodic; this approximation will be good if *w*(*t*) decays to zero away from *t* = 0 sufficiently fast so that ∑_*k* ≤− ⌈*P/*2⌉ +1_ |*a*_*k*_|≈ 0 and ∑_*k*≥ ⌊*P/*2⌋_|*a*_*k*_|≈ 0. This change of boundary conditions gives rise to a linear system of ordinary differential equations with a circulant coefficient matrix that we call *A*:

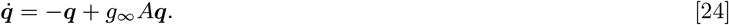

The following analysis simplifies if we reindex the coefficients as *a*_*k*_ for *k* ∈ {− ⌊*P/*2⌋, − ⌊*P/*2⌋+1, …, ⌊*P/*2⌋}. For even *P* this will leave us with *P* + 1 coefficients (one more than is contained in *A*), so in this case we require *a*_− ⌊*P/*2⌋_ = 0. The periodic boundary conditions (in *μ*) imply that *A* is circulant, which means that *A* can be diagonalized by Fourier modes, with eigenvalues that take a simple form. Specifically, *A* being circulant means that − *I* + *g*_∞_*A* has eigenvalues 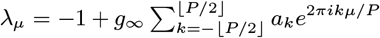.

Since the network dynamics are bounded by the saturating nonlinearity *ϕ*, overlaps (*qμ t*) will remain finite over time, which implies that max_*μ*_ **ℜ***λ*_*μ*_ ≤ 0. To be stable, the solutions must also not decay to zero, which implies that max_*μ*_ **ℜ***λ*_*μ*_ ≥ 0. Hence stable solutions of the system with periodic boundary conditions satisfy max_*μ*_ **ℜ***λ*_*μ*_ = 0. We next relate the condition max_*μ*_ **ℜ***λ*_*μ*_ = 0 to the coefficients *a*_*k*_. We first note that

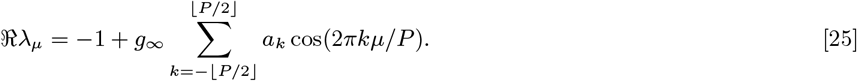

Each of the cos factors is maximized at *μ* = 0; however, some of the *a*_*k*_ may be negative, which makes the location of the maximum for the sum less clear. Hence, we pair the terms in Eq. (25) as follows:

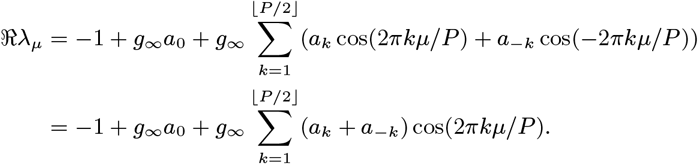

(For *P* → ∞, this rearrangement is valid provided the sum is absolutely convergent). We next assume that each of the forward terms *a*_*k*_ is positive and larger in magnitude than the corresponding backward term *a*_− *k*_. Under this assumption, each of the terms in the above sum is positive, so that the maximum occurs at *μ* = 0. Note that this assumption is stronger than necessary; the maximum will occur at *μ* = 0 provided the positive *a*_*k*_ + *a*_− *k*_ terms significantly outweigh the negative. This is a reasonable assumption for getting forward-propagating sequential behavior. Hence we can write

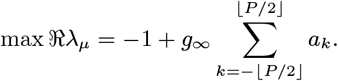

Hence max **ℜ** *λ*_*μ*_ = 0 when Eq. (22) holds (using again the assumption that the terms *a*_*k*_ for *k* >⌊ *P/*2⌋ and *k* < ⌊*P/*2⌋ sum to zero).

In summary, we found that stable dynamics implies Eq. (22) under the following assumptions: (1) the limit *g*_∞_ = lim_*t*→∞_ *g*(*t*) exists, (2) ∑_*k*≤− ⌈*P/*2⌉ +1_|*a*_*k*_| ≈ 0 and ∑_*k*≥ ⌊*P/*2⌋_|*a*_*k*_| ≈ 0 (so that a periodic version of the nonperiodic system can be fined), and (3) the forward terms *a*_*k*_ for positive *k* sufficiently dominate the backward terms *a*_− *k*_, so that the maximum of 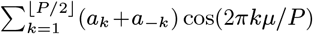 occurs at *μ* = 0. If taking *P* → ∞, we additionally require the series 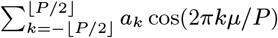 to be absolutely convergent so that the terms can be rearranged.

When will the network dynamics satisfy the stability condition of Eq. (22)? Since *g*(*t*) is bounded from above by *G*(*ρ*^2^) (see Appendix 1), clearly the stability condition cannot be satisfied if 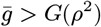; in this case the network activity will decay to zero. What if 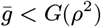? We can reason that the network will indeed satisfy the stability conditions. This is because, if the network activity is vanishing, then the argument ∥*h*(*t*) ∥^2^ + *ρ*^2^ to *G* will approach *ρ*^2^. This causes *g*(*t*) = *G*(∥*h*(*t*) ∥^2^ + *ρ*^2^) to grow to a value where it causes the network activity to grow (since *G*(*ρ*^2^) is larger than the critical boundary of stability of 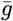). A similar line of reasoning can be found in (37).

#### Bidirectional connectivity – explicit solution

Here we derive an expression for a solution to the mean-field system with bidirectional connectivity. We include this derivation for completeness and to illustrate the difficulties that arise in deriving a simple exact expression for the solution. The reader may prefer to skip to the next sections showing our methods for finding approximate solutions. We start with the system of equations

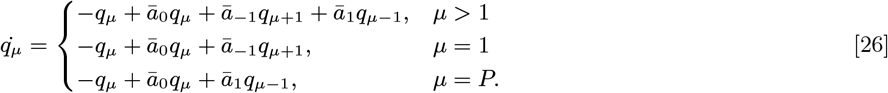

In this case, the coefficient matrix *A* for the system written as 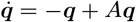 has eigenvalues

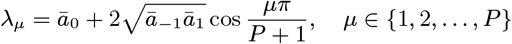

and eigenvectors ***v***^*μ*^:

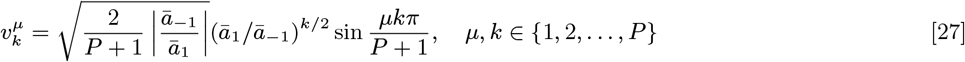

The solution to Eq. (26) is then

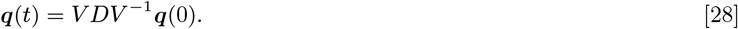

where the columns of *V* are the eigenvectors ***v***^*μ*^ and where *D* is a diagonal matrix with 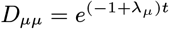. Letting *V* ^*^ denote the conjugate transpose of *V*, we see that *V V* ^*^ is a diagonal matrix with the first entry being equal to 1. To see this, an easy calculation shows that

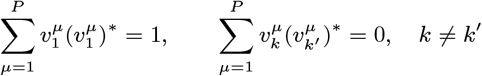

where the asterisk denotes the complex conjugate. Hence, if we use the initial conditions *q*_*μ*_(0) = *δ*(*μ* − 1), we have that 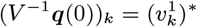. Hence we can write Eq. (28) as

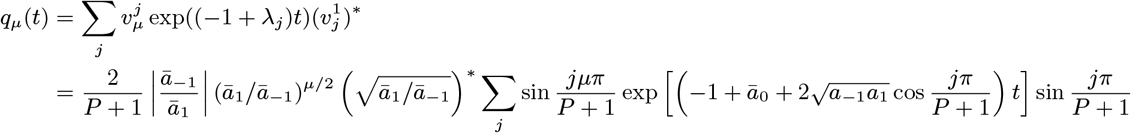

Taking *P* → ∞ this becomes an integral

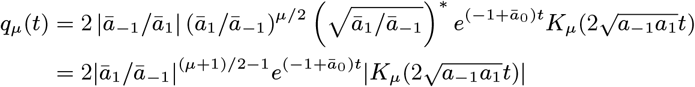

where

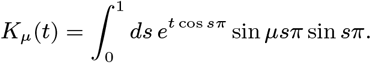

Noting that

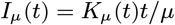

is the modified Bessel function of the first kind, this equation can be written

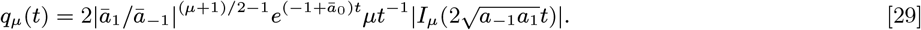

Instead of analyzing these equations further, we proceed with approximate methods that will generalize to more cases (i.e. more nonzero coefficients *a*_*k*_).

#### Bidirectional connectivity – periodic boundary conditions

To further analyze the behavior, we convert the boundary conditions of Eq. (26) to periodic. This yields the following system of equations

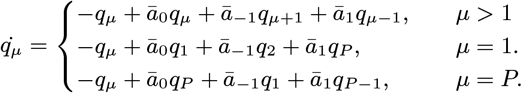

Note that the periodic equations may not match the solutions to the nonperiodic equations exactly, even as *P* → ∞. This makes our simulations important for verifying that the approximations are useful.

The periodic equations can be written

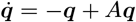

where

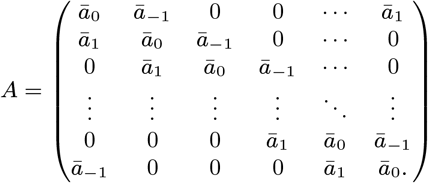

Since is circulant, it can be diagonalized by Fourier modes. Specifically, the *μ*th eigenvalue and eigenvector of + can be written 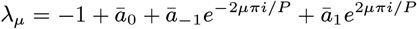 and 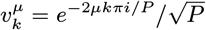. Using the initial condition *q*_*μ*_(0) = *δ*(*μ* − 1), the solution can then be written

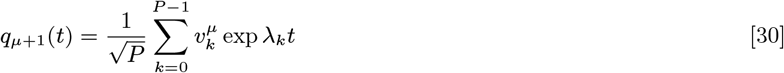

which becomes for *P* → ∞

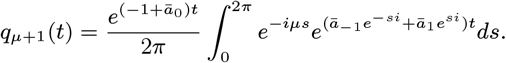

Note that in taking *P*→ ∞, we still require that *N* is much larger than *P* so that the mean-field equations are valid. Simplifying the integrand results in

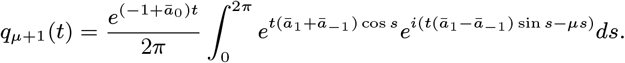

and using the 2*π*-periodicity of the integrand

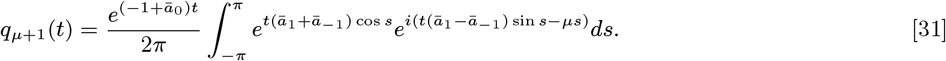

We next have some choice of method for approximating this integral. A simple approach would involve taking a truncated Taylor expansion of the trigonometric functions in the integrand, with the hope that this resolves into an integral that can be evaluated. A more sophisticated and principled approach would be to use a saddle point approximation, where *t* is treated as a variable that is becoming large. We will start with saddle point approximations and show the limitations of this approach in this instance. We will then compare the result with the naive Taylor expansion approach. To get the essential information, the reader may wish to skip to this Taylor expansion approach.

#### Bidirectional connectivity – saddle point approximation

We proceed by performing a saddle point approximation of the integral in Eq. (31). To do so, we write the integral as

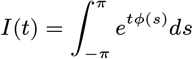

where *ϕ*(*s*) = *β* cos *s* + *iα* sin *s − γs* and *β* = *ā*_1_ + *ā*_− 1_ and *α* = *ā*_1_ − *ā*_− 1_ and *γ* = *μ/t*. Note that we are treating *μ/t* as a finite scalar, so *μ* and *t* are growing large together.

For the saddle point approximation, we first extend *ϕ* to a function over the complex numbers and find the critical points of *ϕ* (**ℜ***ϕ* are saddle points at these critical points in the complex plane). When *γ* is large enough, there are two saddle points that both lie on the imaginary axis. To find these, we can take *s* = *iy* and find the roots of *ϕ*^′^(*iy*). Letting *u* = *e*^*y*^, *ϕ*^′^(*iy*) can be written *ϕ*^′^(*iy*) = − *iβ*(*u* − 1*/u*)*/*2 + *iα*(*u* + 1*/u*)*/*2 − *iγ*. The roots of the resulting quadratic are

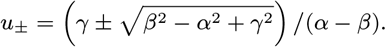

Here we see that the condition that these roots exist is that *γ*^2^≥ *β*^2^ − *α*^2^ since *u* must be real. We next need to calculate the angle of approach for these saddle points. This is given by arg *ϕ*^′′^ evaluated at the saddle points. This shows that the angles are − *π/*2 and 0 for the saddle points in the *y* > 0 positive half-space and *y* < 0 negative half-space, respectively. The contribution of the saddle point in the positive half-space is pure imaginary, so we can focus on the saddle point in the negative half space, which we denote *s*^*^. Given the negative root *u*_−_ above, it is straightforward to compute that

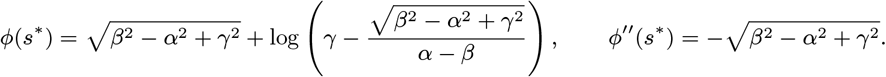

The resulting saddle point approximation yields

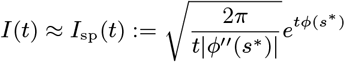

so that

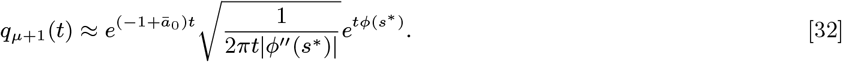

To find the peak times, we seek the roots of 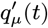. Given the saddle point approximation above, it is straightforward to compute 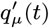. This expression, even in the asymptotic limit *t*→ ∞ (with *μ/t* constant), does not appear to have roots that can be expressed in an explicit equation for *t*.

Given that the roots of the saddle point approximation are unhelpfully complex expressions, we next try taking a saddle point approximation of 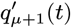; i.e., instead of finding a saddle point approximation and then taking a derivative, we first take a derivative and then find the saddle point approximation. The derivative 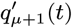 is

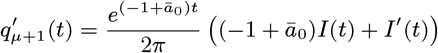

where

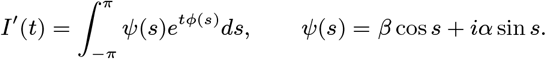

Taking a saddle point approximation of this involves taking a saddle point approximation of *I*(*t*) and *I*^′^(*t*); we have already done the former, and the latter is simply

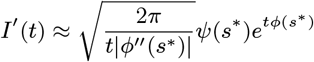

**Fig. S2.**
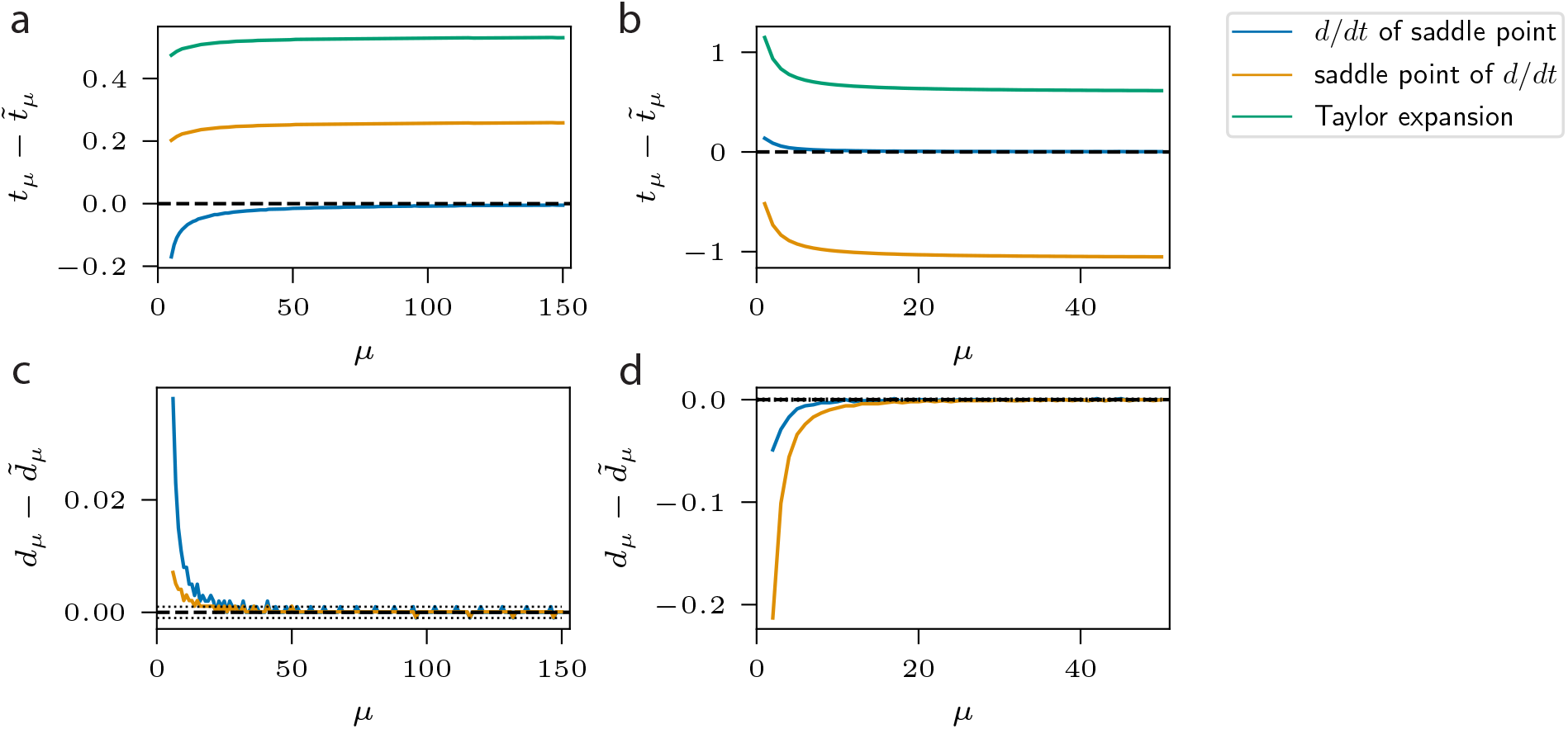
Comparing approximations of *t*_*μ*_ to true peak times. In these plots, the difference between the true peak times *t*_*μ*_ and approximations 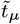 is plotted across *μ*. Blue line, orange line, and green line correspond to the peaks of the saddle point approximation of *q*_*μ*_(*t*), the roots of the saddle point approximation of 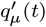, and the peaks of the Taylor expansion approximation of *q*_*μ*_(*t*), respectively. The python `scipy.signal` utility `find_peaks` is used to find the peak times numerically for the blue line. True peak times (dashed lines) are found by numerical quadrature of the integral expression in Eq. (19), followed by using the `find_peaks`. **a**) Coefficient values *a*_− 1_ = *−* .3, *a*_0_ = .1, and *a*_1_ = 1.8. **b**) Coefficient values *a* _1_ = .2, *a*_0_ = .2, and *a*_1_ = .8. **c**)-**d**) Difference of peak differences *d* = *t − t* _1_ with approximations 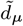. Note that the orange and green lines coincide. Dotted lines denote the precision ceiling due to the step size *dt* = 0.001. **c**) Coefficient values *a*_− 1_ = *−* .3, *a*_0_ = .1, and *a*_1_ = 1.8. **d**) Coefficient values *a*_− 1_ = .2, *a*_0_ = .2, and *a*_1_ = .8.

where *s*^*^ is the critical point of *ϕ* as above. Hence

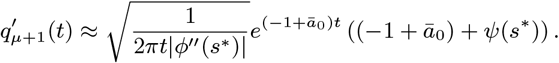

We again look for the peak times by finding the roots of this expression, which occur at ((−1 + *ā*_0_) + *ψ*(*s*\*)) = 0. Evaluating *ψ*(*s*\*) is similar to evaluating ^′′^(^*^) and yields 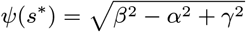. Substituting *β* = *ā* + *ā*, α = *ā*_*1*_ + *ā*_*−1*_, *α* = *ā*_1_ − *ā*_*−1*_, *တ*_*1*_ = 1 − *ā*_*1*_ − *ā*_*−1*_ and *γ* = *μ/t* into this expression and solving 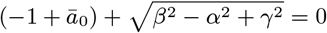 for *t* yields

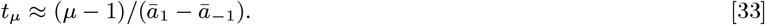

This is the same as Eq. (12) up to a constant offset.

While this is a fair approximation, the differences between this estimate and the true peak times typically remain non-vanishing even as *t* and *μ* approach infinity; see the blue and orange lines of Figs. S2a and S2b. However, the differences between the true peak differences *d*_*μ*_ = *t*_*μ*_ − *t*_*μ* − 1_ and this estimate do vanish with increasing *t* and *μ*; see the blue and orange lines of Figs. S2c and S2d.

Considering that the saddle point method yields either (1) an estimate for *t*_*μ*_ that doesn’t appear to have a simple closed-form expression, or (2) an estimate that is not asymptotically precise, we seek a simpler approach. This simpler approach derived in the next section has the additional advantage of generalizing to the case of arbitrarily many nonzero coefficients *a*_*k*_.

#### Bidirectional connectivity – Taylor expansion

In our next approach to approximating the integral Eq. (31), we expand the trigonometric functions around *s* = 0 to get

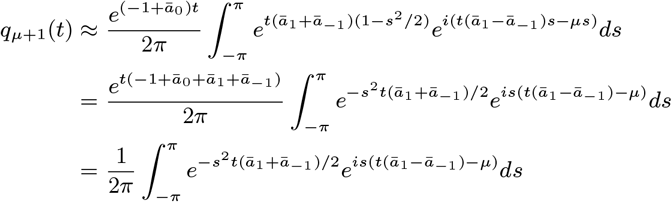

where in the last step we have used that ∑*ā*_*k*_ = 1. Letting *α* := *ā*_1_ − *ā*_− 1_ and *β* := *ā*_1_ + *ā*_− 1_, we rewrite the integral as

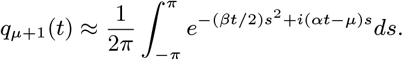

While we can write this integral in terms of error functions, a simpler expression results if we are able to integrate over the whole real line. We note that across a range of reasonable choices of the *a*_*k*_, the integrand is vanishingly small outside of the interval [− *π, π*]. Hence we can indeed approximate the integral with one over the real line. This results in

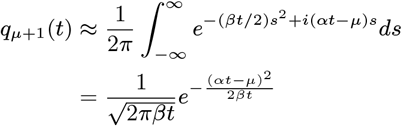

The locations *t*_*μ*_ of the peaks are given by the positive root of 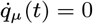, which when using the approximation above yields

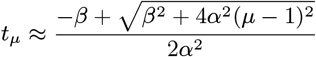

Taking the asymptotic limit of large *μ* and taking the positive root yields Eq. (14):

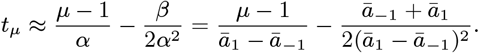

Similar to the estimate yielded by taking a saddle point approximation of 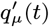, this estimate is not asymptotically exact; see the green lines in Figs. S2a and S2b. Depending on the choice of *a*_*k*_, the approximation can be bettor or worse than the saddle point approximation Eq. (33); compare Fig. S2a with Fig. S2b. However, as before the differences *d*_*μ*_ := *t*_*μ*_− *t*_*μ* − 1_ are asymptotically correct; see Figs. S2c and S2d. Hence we can consider this approximation to be practically at least as useful as that yielded by taking saddle point approximations. An extra benefit of this approach is that it generalizes; see the next section.

#### Generalizing the computation to shift-invariant connectivity

The above technique of using Taylor expansions to find an approximate solution for *q*_*μ*_ extends to the general case of Eq. (10) with periodic boundary conditions in *μ*. As in the derivation of 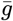 above, we index the coefficients as *a*_*k*_ for *k* ∈ { *P/*2⌋, − ⌊*P/*2⌋ + 1, …, ⌊*P/*2⌋}. Then the coefficient matrix *A* is circulant, and the eigenvalues of *I* − *A* are 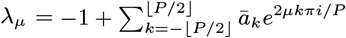. Then grouping terms and taking the limit as *P* → ∞ results in

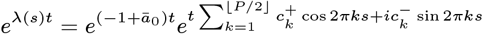

where 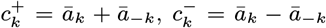, and *s* ∈ [0, 1]. Hence, using the eigenvectors Eq. (27) of circulant matrices in the expression Eq. (30) (which is also valid here),

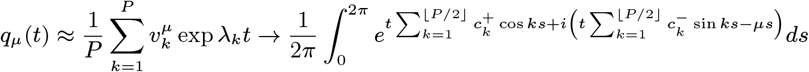

as *P* → ∞. Redefining the bounds of integration to [− *π, π*] and expanding the trigonometric functions around *s* = 0 as above yields

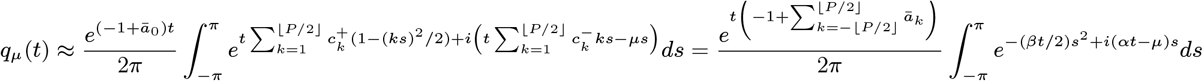

where 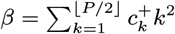 and 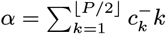. Again extending this to an integral over the real line as before, as well as noting 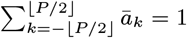, yields the solution

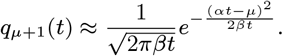

As before, the positive root of 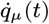 is asymptotically

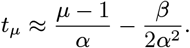

which is Eq. (12). This approximation will be used for the remainder of the Appendices.

#### Two forward terms

In the case of two forward terms, the system of equations is

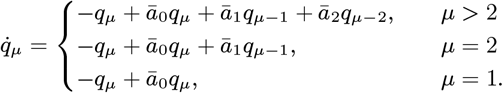

Using Eq. (12) yields the approximate peak times of Eq. (15):

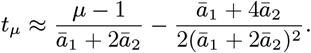

#### Exponential kernels

Here we derive the peak times *t*_*μ*_ for exponential kernels of the form 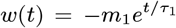 for *t* < 0 and 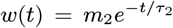 otherwise. As usual, we suppose that the coefficients *a*_*k*_ are indexed as *k* ∈ {− *P* ^′^, − *P* ^′^ + 1, …, *P* ^′^] for some integer *P* ^′^ and define *a*_*k*_ = 0 for *k* > *P/*2 if necessary. Given the approximation Eq. (12), we need only compute 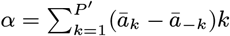 and 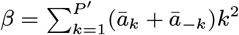. To start we note that the relationship between the coefficients *a*_*k*_ and the kernel *w* takes the form

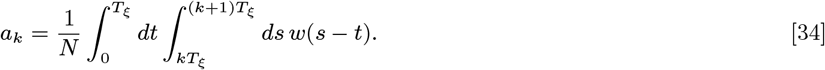

Then, for *k* > 0,

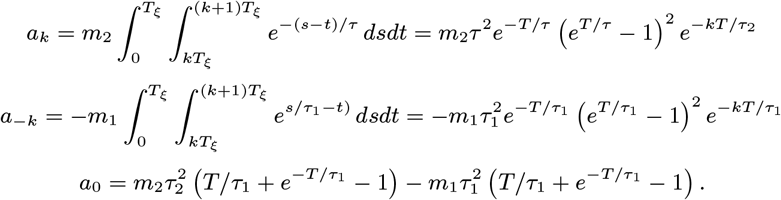

From this we compute

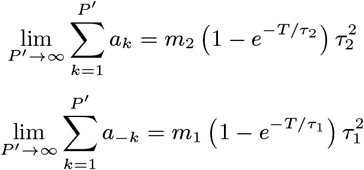

Hence, for large *P* ^′^,

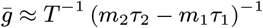

Similarly

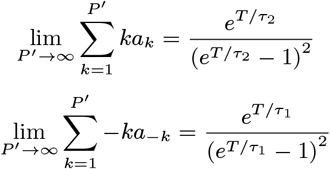

Hence, for large *P* ^′^,

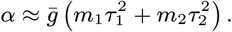

A similar calculation yields

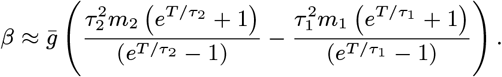

## Appendix 3

Here we give the details of the network simulations. Differential equations are solved with forward Euler integration with a timestep of *dt* = 0.075, except for Fig. 1 which uses *dt* = .0375 and Fig. 3e, which uses *dt* = 1*/*15. Recall that the length of the input patterns is denoted by *N* and the number of patterns in the sequence is *P*. In Figs. 1 and 3 to 5 and Fig. S2, *N* = 35000. In Figs. 6 and 7, *N* = 40000. In Fig. 1, *P* = 40. In Figs. 3 to 5, 6b, 6c and 7 and Fig. S2, *P* = 100. In Fig. 6d, *P* = 60. In Figs. 2 to 6 and Fig. S2, *P* = 100.

Throughout, we use the nonlinearity Eq. (18) with *r*_span_ = 2, *r*_center_ = 0, *θ* = 0, and *σ* = 0.1.

Error bars (shaded regions) are computed automatically by the Seaborn plotting library, which uses bootstrapping to compute 95% confidence intervals.

## Footnotes

* We can also define the sequence state represented at a time *t* by arg max_*μ*_(*q*_*μ*_(*t*)), but the *t*_*μ*_ are more mathematically convenient objects to consider.

† Note the contrast to the dynamical systems literature, where stability typically means that the solutions remain finite.

